# Indoril: An I-PV Add-On for Visualization of Point Mutations on 3D Cartesian Coordinates

**DOI:** 10.1101/148122

**Authors:** Ibrahim Tanyalcin, Julien Ferte, Taushif Khan, Carla Al Assaf

## Abstract

**Summary:** One of the main goals of proteomics is to understand how point mutations impact on the protein structure. Visualization and clustering of point mutations on user-defined 3 dimensional space can allow researchers to have new insights and hypothesis about the mutation’s mechanism of action.

**Availability and Implementation:** We have developed an interactive I-PV add-on called INDORIL to visualize point mutations. Indoril can be downloaded from http://www.i-pv.org.

**Contact:** ║

**Supplementary Information:** Please refer to the supplementary section and http://www.i-pv.org.

## 1 INTRODUCTION

It is often of interest for researchers to understand how point mutations (SNVs – *single nucleotide variations*) affect the protein structure. There are several programs that utilize sequence and/or structure information to predict the mutation effect such as POLYPHEN2 (Adzhubei et al. 2010), SIFT (Kumar et al. 2009). Other programs such as I-TASSER (Yang et al. 2015) work on 3D structures modeled from raw fasta sequences. At some level, integration of all aforementioned programs can be useful in understanding mutation effect. For instance, the position of amino acids in 3D space can be combined with sequence based predictor scores and visualized. Unfortunately, neither sequence related information nor structural data alone yields completely accurate results and researchers often have to resort to comparison between different algorithms. It also useful to retain chemical property of amino acids and other user generated data during this comparison where researchers can quickly navigate back and forth between different settings. Therefore, we have developed an I-PV (Tanyalcin et al. 2015) add-on called INDORIL to visualize point mutations in user supplied variation/vcf files. All elements rendered in INDORIL are interactive with mouse-over explanations and clickable buttons. It allows selective display/hiding of SNVs based on several parameters. INDORIL can be downloaded from http://www.i-pv.org.

## 2 METHODS

INDORIL is built on top of I-PV, therefore when the user generates an html output using I-PV (sequence in fasta format, variation file in tsv or vcf format and conservation scores in plain text, Tanyalcin et al. 2015), all the necessary data to run Indoril are either pre-stored in JSON format or pulled-out from the DOM (Document Object Model). The user only needs to click on the INDORIL icon in the lower left corner of the hmtl output. Several functions can be accessed using the ‘Show Functions’ options in the INDORIL window as also shown in figure 1.

**Fig. 1.**
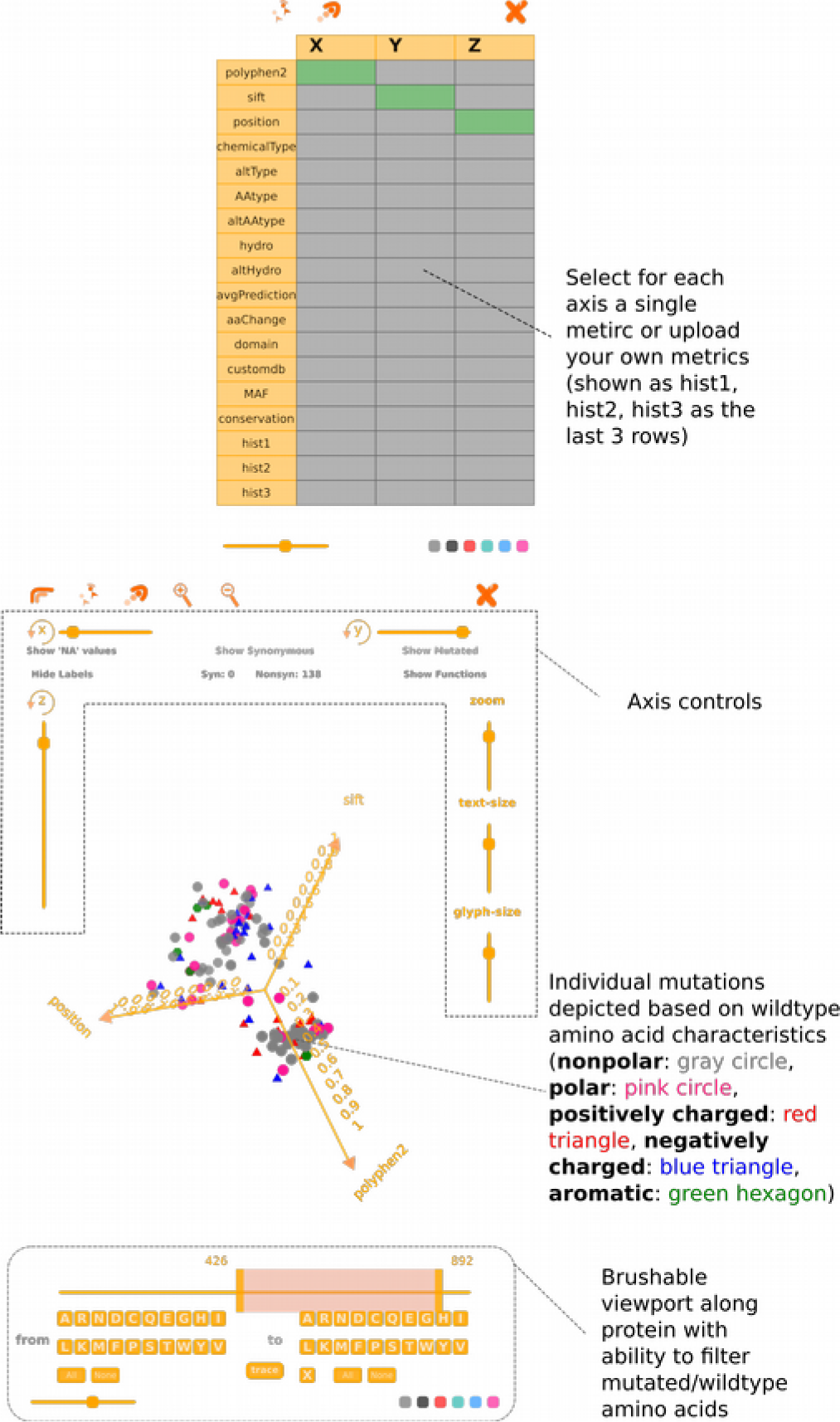
Indoril overview. The add-on is an extension of I-PV, therefore no additional files are needed to initiate an INDORIL instance. The user can open up a new window by clicking on the lower left icon in default I-PV output. Clicking on the “Show Functions” button on the upper right corner brings the list of different functions the user can choose from.

Written in vanilla Javascript and partially D3 v3.5.17 (Bostock et al. 2011), INDORIL works with all modern browsers. Performance of Chrome is the highest due to the use of V8 Javascript engine. Depending on the hardware and the number of mutations to be displayed, framerate drops can be observed in Firefox due to single CPU core restrictions.

## 3 RESULTS

In this article we demonstrate the graphs generated by I-PV for P53 (ENST00000269305), TNF-α (ENST00000449264) and NfκB1 (ENST00000505458). The data necessary to generate the html files are available from Ensembl Biomart utility (http://www.ensembl.org/biomart/, Kinsella et al. 2011) and also contained within the software package.

P53 (OMIM: 191170) is one of the main tumor suppressors genes that regulate cell cycle. Its activity is often reduced in cancer cells by either mutations on itself or mutations affecting other members of the same signaling pathway (Vousden et al. 2007). Figure 2 i shows 4 of the hotspot mutations that occur at position 175. The x, y and z axes show the average of SIFT and POLYPHEN2 prediction scores, conservation and mutated amino acids (abbreviated as altAAtype) respectively within a lattice of 1671 mutations along the entire length of the protein. The 4 mutations (p.R175S, p.R175P, p.R175L and p.R175H) are linked with red lines to better illustrate how they are related to one another with the chosen set of functions.

**Fig. 2.**
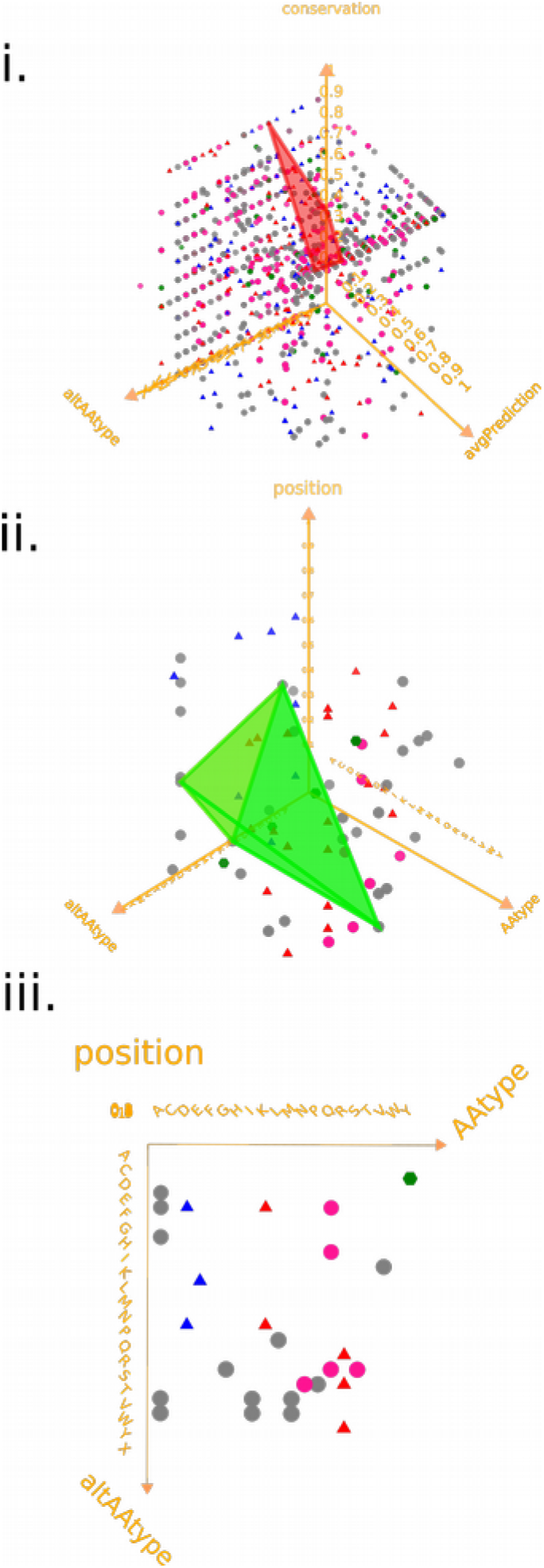
Indoril use cases. **(i)** Construction of mutation lattice and linking of particular SNVs. Symbols are used in the same context as in I-PV main framework (red triangle: positively charged, blue triangle: negatively charged, gray circles: non-polar, purple/pink circles: polar, green hexagon: aromatic) **(ii)** Construction of surfaces with chosen mutations. **(iii)** Construction of basic mutation matrix display (x axis: wildtype amino acid letter, y axis: mutated amino acid letter). Additionally, users can upload their custom data (simple text file with one column of real numbers) and sort/link/cluster mutations based on this custom data.

TNF-alpha is an proinflammatory cytokine that is mainly secreted by the cells of the innate immune system. Apart from suppressing tumor growth, it is detrimental in excess and is involved in autoimmune disorders (Ruuls et al. 1999). Figure 2 ii shows 4 point mutations involving nonpolar amino acids (p.A94T, p.V10M, p.G57S and p.I194N) that have relatively high SIFT and POLYPHEN2 scores. These mutations are linked and re-rendered with x, y and z axes showing original amino acid letter, position along protein and mutated amino acid letter respectively. The links can be drawn to render closed objects depicting volumes as shown in this example.

NfκB1 is a ubiquitous transcription factor that is activated as a result of various cytokines and other extra or intra cellular stimuli. It can lead to activation of inflammatory pathways whereas its constant suppression can result in apoptosis, inappropriate cell growth or cellular growth delay (**Chen et al.**). Figure 2 iii shows how the user can generate a conventional mutation matrix plot where x axis shows the 20 possible wildtype residues (abbreviated as AAtype) and the z axis shows the 20 possible amino acids that can take the place of the wildtype residues. The y axis can be any of the available functions. In Figure 2 iii, the rotation around x axis is used to conceal y values. The user can also remove the diagonal of synonymous mutations from the matrix by clicking on the “Hide synonymous” mutations button.

One of the main problems in visualization of genetic information is integration of different databases. Since INDORIL is built on top of I-PV, the user does not need to provide additional info. On the other hand, if the user has scores from variety of different tools, their scores can be easily uploaded to I-PV (hist1, hist2 or hist3 options shown in Figure 1). After checking the integrity of the uploaded files, three such set of scores can be active at a time. The function panel allows the user to apply different algorithms and see their effects on clustering of mutations. Any linked mutations and rendered polygons at the time of selecting functions are automatically updated. Like in I-PV, the user can extract any frame of the visualization tools as an svg file for a high quality publication ready image.

## 4 DISCUSSION

One of the main challenges in genomics and proteomics is the identification of a given variant is disease causing or not; this has been a central issue since the completion of human genome project (Landen et al. 2001). Several approaches have been implemented for assessing point mutations, for exomiser Exomiser which utilizes various databases to filter vcf files (Robinson et al. 2014). Other approaches are based on machine learning algorithms that include, but are not limited to variant information (Kircher et al. 2014; Maxwell et al. 2015). One limitation of such approaches is that visualization and analysis takes place at different levels.

During the decision process, it can be useful for researchers to rank, visualize and cluster the point mutations in 3d space where the axes can show the values of either user defined functions or built-in features. Especially in monogenic disorders such visualization of SNVs can reveal distinct clusters and landscapes which otherwise would be easily missed with conventional visualization techniques. Visualization of mutations as in Figure 2 takes advantage of human visual cortex as well as allowing researchers to edit the underlying data representation. As opposed to visualization, inferring distances between a pair of mutations from a simple table of n mutations would result in n(n1)/2 comparisons before any verdict can be given.

For example, a user can assign different metrics to each axis such as, hydrophobicity, chemical property, amino acid type, minor allele frequency and custom user generated meta data (automatically normalized between 0-1). These values are then mapped onto the mutation data based on their residue number, and the user can link different mutations to each other (eg. mutations that result in a similar phenotype) while quickly switch back and forth between different configurations. If the said mutations cluster under a certain configuration, this suggests that those metrics might be a possible surrogate for predicting mutation effect. Such insights can further direct researchers to form plausible hypothesis about the mutation’s mechanism of action (such as a polar to aromatic amino acid change at certain regions/domains with polyphen/sift/predictor score above a certain value). Especially in areas where multiple mutations need to be analyzed with vague genotype-phenotype correlations, we believe our tool can give researchers an edge about getting the best out of their data sets.

Having an extra layer of visualization concurrent with analysis (eg. chosing different functions and instantly receiving feedback on each mutation) can help researchers gain more control on the variants they wish to focus on. With our tool, researchers can easily isolate selected variants, link them and see if these variants show any clustering preference under different scoring. In a typical layout with around 400 mutations and using various in-built filters, mutation numbers can be brought down to levels feasible to be assessed by human eye. For larger datasets, the function that handles data to the INDORIL renderer can be called manually, which exposes the data object passed to INDORIL. This object can be stringified and exposed to 3^rd^ party clustering and analysis applications. Such features are not possible in conventional state of the art feature viewers which adopt linear sliding window formats. The functions implemented in INDORIL are elementary and all selected functions return real number values between 0 and 1 which also makes it easy to superimpose results between different proteins for comparison. INDORIL can be downloaded from https://github.com/IbrahimTanyalcin/I-PV.

## ACKNOWLEDGEMENTS

IT wrote the main text and prepared figures. JF and CA tested the software. JF, TK, CA, Wim Vranken and Marianne Rooman reviewed the manuscript, added suggestions and gave feedback. JF and TK helped poster presentation at ECCB 2016 (poster number P_Pr043). CA designed the layout of the software.

